# McsB regulates CtsR thermosensing through periphery arginine phosphorylation

**DOI:** 10.1101/2024.12.26.630380

**Authors:** Huahuan Cai, Boyang Hua, Jie Hu, Yiben Fu, Xinlei Ding, Gege Duan, Yeting Guo, Xing-Hua Xia, Yufen Zhao

## Abstract

When cells sense an elevated temperature in the environment, the bacterial master transcription repressor CtsR becomes phosphorylated and inactivated by the arginine kinase McsB to initiate the expression of heat-shock genes. Here, we utilize a fluorescence intensity shift assay (FISA) based on the protein-induced fluorescence enhancement (PIFE) effect to monitor the DNA-CtsR-McsB interactions in real time. Our single-molecule analysis reveals that CtsR binds rapidly and stably to the cognate DNA, and that McsB is able to transiently interact with CtsR *in situ* of the target DNA. We determine the binding kinetics between McsB and the DNA-bound CtsR by single-molecule real-time binding assays, with *k_on_* and *k_off_* of 0.75 μM^-1^ s^-1^ and 0.34 s^-1^, respectively. This interaction with McsB does not remove CtsR from the DNA, but lowers the temperature threshold for CtsR dissociation and alters its thermosensing behavior. Mass spectrometry, mutational analysis and structural simulation results together suggest that the phosphorylation of several periphery arginine residues on CtsR, which reduces the binding energy of the CtsR-DNA interaction, underlies a plausible molecular mechanism for this effect. Taken together, these results provide insights into how McsB regulates the CtsR-DNA complex and highlight the functional importance of CtsR periphery arginine residues in the bacterial heat-shock response.

## Introduction

First discovered in 1976, arginine phosphorylation is a widespread and crucial PTM in both prokaryotic and eukaryotic cells.^1,2^ It is estimated that over 10,000 phosphorylated arginine sites exist in the bacterial and human proteomes, showcasing the scale and prevalence of this form of N-phosphorylation.^3–7^ Arginine phosphorylation on a protein can influence its structure, activity, stability, interactions and cellular localizations. In bacteria, numerous important biological processes are influenced by this PTM, such as gene expression regulation^8,9^ and cell division,^10^ highlighting its broad roles in bacterial adaptation, persistence, and virulence.^11–15^ Despite the importance of this PTM, the identity of the kinase responsible for phosphorylating arginine residues remained elusive for a long time.

McsB is the founding member of the cellular arginine kinases and is shown to phosphorylate several arginine residues on the bacterial master transcription repressor CtsR.^9^ Recent structural studies reveal that McsB has an N-terminal phosphagen kinase (PhK)-like phosphotransferase domain and a C-terminal dimerization domain containing a phospho-arginine-binding pocket.^16^ McsB typically functions as a multimer, ranging from dimers in *Geobacillus stearothermophilus* to mostly higher-order oligomers in *Bacillus subtilis*.^17^ The state of oligomerization is thought to regulate the kinase activity of McsB, since mutations that disrupt the dimer formation markedly reduce the activity.^16^ McsB autophosphorylation is important to its function, as the dimerization process is mediated through the phospho-arginine residues.^16,17^ Besides the autophosphorylation and oligomerization, the kinase activity of McsB from *B. subtilis* also requires the cofactor McsA, which enhances the kinase activity of McsB by reorganizing the catalytic site, accelerating the enzyme’s turnover rate, and increasing its thermostability.^18,19^ In contrast, McsB from *G. stearothermophilus* on its own is highly active *in vitro*, with activity even surpassing the *B. subtilis* McsB in complex with McsA.^18^

The McsB-CtsR system plays an indispensable role in regulating cellular responses to harsh conditions and promotes cell survival under stress situations. For example, in Gram-positive bacteria, the heat stress causes protein misfolding-induced toxicity and triggers protein quality control responses mediated by the transcription repressors HcrA and CtsR.^20–25^ Under normal conditions, CtsR forms a dimer that specifically binds to the promoter region of the *clpC* operon and represses the expression of the operon which includes several heat-shock genes, such as *ctsR*, *mcsA*, *mcsB* and *clpC* in *B. subtilis*.^26^ Based on previous structural studies, the CtsR dimer interacts with DNA mainly through its several arginine residues, with R28 and R49 bind the major groove whereas R62 invades deeply in the minor groove.^9^ Upon heat exposure, a glycine-rich loop in CtsR senses the temperature change and promotes CtsR dissociation from the DNA, thus activating the expression of the *clpC* operon.^27^ Once dissociated, R28, R49 and R62 become exposed and can be phosphorylated by the arginine kinase McsB. Arginine phosphorylation inverses the charges on the arginine residues and prevents its interaction with the negatively charged DNA backbones.^9^ Furthermore, arginine phosphorylation also signals for protein degradation by the ClpCP protease system in the prokaryotes, akin to the function of ubiquitination in the eukaryotes.^28^

In addition to the heat stress, McsB and arginine phosphorylation are also involved in the regulation of CtsR in response to other environment stimuli, such as the oxidative and metal ion stresses.^29–31^ For instance, the oxidative stress can lead to the inactivation of CtsR and de-repression of stress genes in a McsB and arginine phosphorylation-dependent manner. In this case, however, it is not understood how the core arginine residues (*i.e.*, R28, R49 and R62) that make direct interactions with the target DNA become exposed to McsB in the absence of temperature changes. Moreover, apart from the three core arginine residues, it was shown that McsB can phosphorylate five other arginine residues on CtsR (*i.e.*, the periphery arginine residues).^9^ It is unclear if and how the phosphorylation of these arginine residues affects the CtsR-DNA binding affinity.

To address these questions, we employed several single-molecule fluorescence techniques to study how McsB regulates CtsR and its temperature-dependent DNA binding affinity. The toolbelt of single-molecule fluorescence techniques, which commonly include colocalization,^32^ fluorescence resonance energy transfer (FRET)^33,34^ and PIFE^35,36^ methods, are now routinely applied in biological studies. Due to the ability to reveal heterogeneous subpopulations and hidden intermediate states, these tools are ideal for studying the structural and interaction dynamics of biomolecule complexes. In this work, our single-molecule results suggest that McsB transiently but specifically interacts with the DNA-bound CtsR. Although not directly resulting in the dissociation of CtsR from its target DNA, this interaction leads to the phosphorylation of several periphery arginine residues on CtsR and renders the CtsR-DNA complex more sensitive to heat. Our findings further elaborate the function of arginine phosphorylation on CtsR and suggest a hierarchical phosphorylation and regulation mechanism by which McsB loosens the CtsR-DNA complex from the periphery and “works its way” toward the core to achieve the full inactivation of CtsR.

## Results

First, to study the interaction between CtsR from *G. stearothermophilus* and its cognate DNA, we designed a fluorescently labeled DNA probe that exhibited the PIFE effect^36^ upon CtsR binding. Briefly, a fluorophore is placed adjacent to the CtsR consensus sequence so that CtsR binding positions the protein in close vicinity to the fluorophore, resulting in a fluorescence enhancement **(Figure 1A-B and Supplementary Figure 1A)**. The fluorophore used and its labeling position on the DNA were optimized, as these factors were shown to affect CtsR binding and the PIFE effect **(Supplementary Figure 1B)**. The DNA probe is relatively short and holds only binding site for one CtsR dimer. We used this DNA probe and a sequence-scrambled control to determine the binding affinity of CtsR to its target DNA. For the cognate DNA probe, as the CtsR concentration increased, the intensity peak position shifted from a lower value (peak 1 at 385 a.u.) of the DNA probe alone to a higher one (peak 2 at 600 a.u.) at 50 nM CtsR **(Supplementary Figure 1C)**. In contrast, such intensity peak shift was not observed with the non-cognate DNA probe **(Supplementary Figure 1C)**. The y-axis value ratio at these two peak positions was used as a metrics to approximate the CtsR-bound fraction **(Figure 1C)**. Through fitting the curve with the Hill equation, the *K_D_* and n were around 12 nM and 2.1 for the cognate DNA, respectively, whereas the non-cognate control was barely bound by CtsR even at elevated protein concentrations **(Figure 1C)**. These results demonstrate the high affinity, sequence specificity and cooperativity of the CtsR-DNA interaction, in consistent with the structure where a CtsR dimer binds to the target DNA. The measured *K_D_* is also in agreement with the previous result (*K_D_* ∼25 nM) measured by isothermal titration calorimetry.^9^ Based on this result, 25 nM to 50 nM CtsR was used in most of the following experiments to ensure a saturated CtsR dimer binding of the DNA targets.

**Figure 1.**
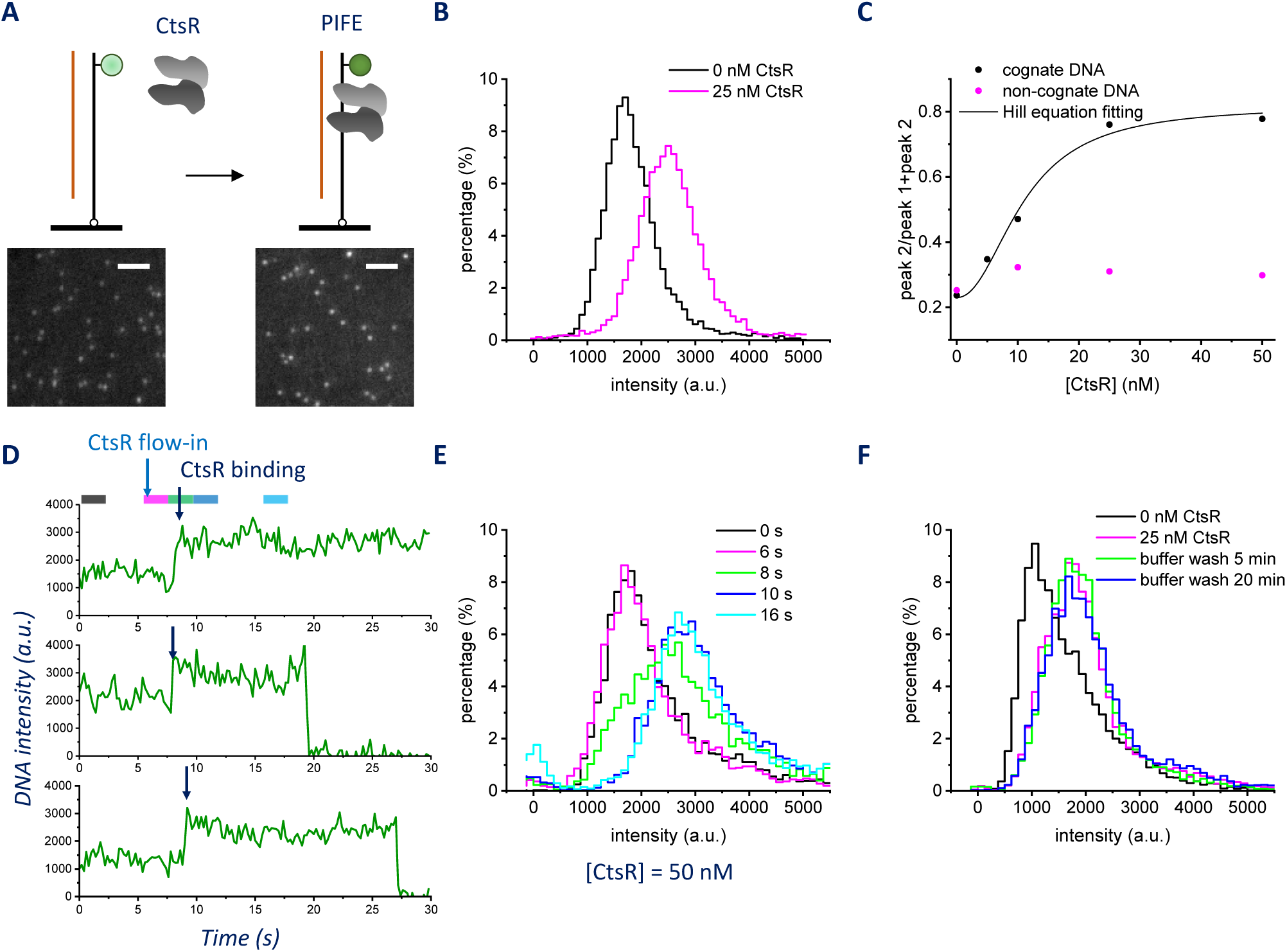
Single-molecule analysis of the CtsR-DNA interaction. (A) Schematic showing the fluorescently labeled DNA probe for monitoring CtsR binding based on the PIFE effect. Inserts are representative images of single DNA probes before and after CtsR binding. The images were displayed at the same contrast with ImageJ. The scale bars correspond to 2.5 μm. (B) Fluorescence intensity histograms of the DNA probe alone (black) and bound by 25 nM CtsR (magenta). Experiments were sequentially performed in the same flow channel. 5-10 random areas containing 1023 and 917 single molecules were imaged to construct the intensity histograms, respectively. (C) The fraction of CtsR-bound DNA as a function of the protein concentration in solution. Black dots represent the cognate DNA, whereas the magenta dots represent the non-cognate DNA. To obtain the *K_D_*, the data were fit with the Hill equation. (D) Example fluorescence intensity trajectories of real-time CtsR binding to the target DNA upon the addition of 50 nM CtsR in the flow cell. The color bars indicate the frames used to construct the intensity histograms of the same color in (E). (E) Time-dependent fluorescence intensity histograms constructed from the acquired trajectories at different time points as indicated by the colored segments in (D). 457 trajectories from the flow-in movie were used to build the histograms. (F) Fluorescence intensity histograms of the CtsR-DNA complex before and at different time points after a buffer wash to remove the excess CtsR in solution. Experiments were sequentially performed in the same flow channel. 5-10 random areas containing 813±82 (mean±standard deviation) single molecules were imaged to construct each intensity histogram.

Next, we explored the feasibility of using the PIFE reporter to monitor CtsR binding in real time. A flow cell was constructed, in which unlabeled CtsR was introduced during live image acquisition **(Figure 1D and Supplementary Figure 1D)**. Before adding CtsR, we detected a stable and low fluorescence intensity level, corresponding to the unbound DNA probe. Upon the addition of CtsR, a single-step increase in the fluorescence intensity was observed, indicating the binding of CtsR to the target DNA. To estimate the binding rate *k_on_*, we constructed a series of time-dependent intensity histograms, which showed that the binding events were largely completed within 4 s after CtsR addition **(Figure 1E and Supplementary Figure 1E)**, indicating CtsR has a rapid DNA binding rate. In most of the single-molecule trajectories, the CtsR-bound high fluorescence intensity was stable until the end of the movie or when photobleaching occurred (*e.g.*, a single-step decrease in the fluorescence intensity to the baseline level, **Figure 1D**). Only a small fraction of the molecules showed a two-step fluorescence intensity decrease which may represent CtsR dissociation (*i.e.*, the loss of PIFE) followed by photobleaching **(Supplementary Figure 1D)**. Furthermore, we did not observe considerable CtsR dissociation after washing away CtsR in solution (**Supplementary Figure 1F**). These results suggest that CtsR has a relatively slow unbinding rate and binds stably to its target DNA. To further assess the unbinding rate beyond the limit of photobleaching, we took intensity histograms 5 min and 20 min after washing away the excess CtsR in solution. At room temperature, CtsR stayed bound to its target DNA for at least 20 min without displaying substantial dissociation **(Figure 1F)**.

McsB phosphorylates several key arginine residues on CtsR, thus reversing the positive charges and impairing its binding to the negatively charged DNA.^9^ To study if McsB can remove the DNA-bound CtsR in our assays, we incubated the CtsR-DNA complex with the autophosphorylated and active McsB (pMcsB) from *G. stearothermophilus*, and took intensity histograms at room temperature 5 min and 20 min after the addition of pMcsB and ATP **(Figure 2A and Supplementary Figure 2A)**. Unlike the free CtsR, which was completely inactivated by pMcsB **(Supplementary Figure 2B-C)**, the DNA-bound CtsR remained active and did not show any loss of PIFE that indicated its removal. Interestingly, the population of the CtsR-DNA complex shifted to slightly higher intensities after the pMcsB and ATP treatment. To rule out the possibility that CtsR was non-specifically stuck to the surface and thus could not be removed by pMcsB, an unlabeled cognate DNA (without biotin) was added in solution to compete with the surface-tethered DNA targets. A substantial down-shift in the intensity histograms was observed, suggesting the successful dissociation of CtsR from the surface **(Supplementary Figure 2D-E)**.

**Figure 2.**
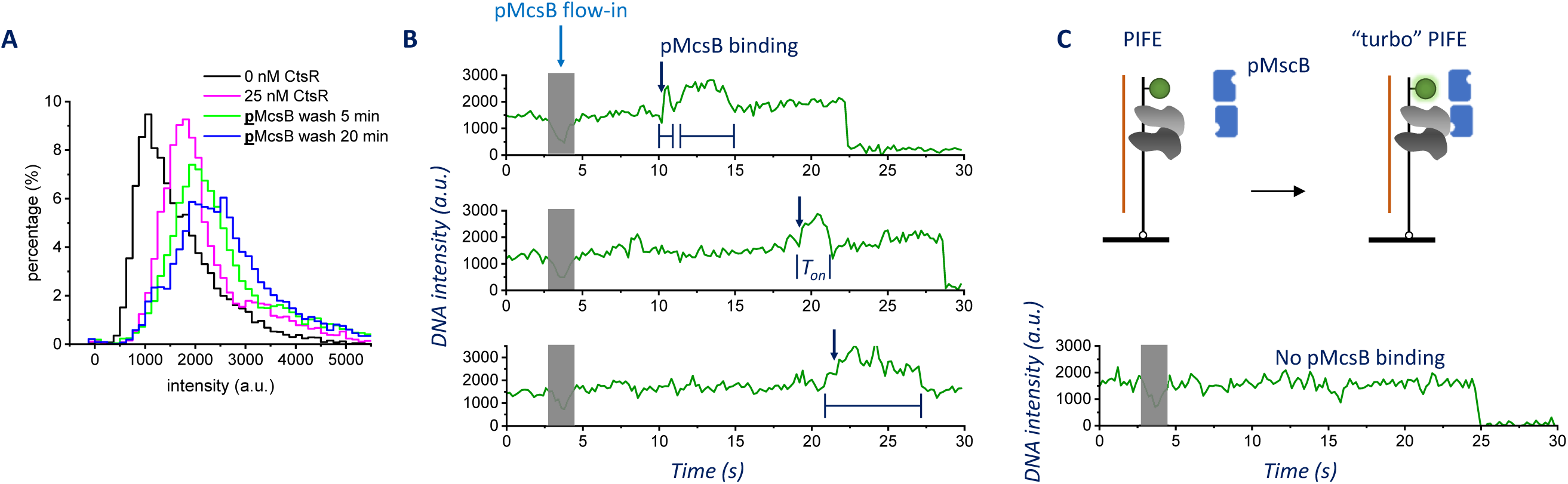
Single-molecule analysis of the pMcsB-CtsR-DNA interaction. (A) Fluorescence intensity histograms of the CtsR-DNA complex before and at different time points after a pMcsB and ATP wash to remove the excess CtsR in solution. Experiments were sequentially performed in the same flow channel. 5-10 random areas containing 899±38 (mean±standard deviation) single molecules were imaged to construct each intensity histogram. (B-C) Example fluorescence intensity trajectories of real-time pMcsB binding to the CtsR-DNA complex upon the addition of 200 nM pMcsB and 0.5 mM ATP in the flow cell. In many trajectories, transient spikes were observed indicating pMcsB binding to the CtsR-DNA complex, while other trajectories did not show such events. The square bracket indicates the pMcsB-bound dwell time. The *k_off_* was estimated to be 0.29 s^-1^ (averaged from 73 T_on_ from 50 single molecules). (C) Schematic showing pMcsB binding to the CtsR-DNA complex as monitored by the “turbo” PIFE effect.

We then performed a real-time flow-in experiment to monitor the CtsR-DNA complex upon the pMcsB and ATP addition **(Figure 2B-C)**. Before the pMcsB flow-in, a stable and high intensity level was observed due to the PIFE effect caused by the bound CtsR. After the pMcsB flow-in, as sometimes marked by an abrupt defocusing and refocusing event, we detected frequent and transient fluorescence intensity increases to an even higher level beyond that induced by CtsR alone in a portion of the trajectories **(Figure 2B)**. Trajectories that displayed such signals were characterized as “dynamic”, which were much less common when a buffer without pMcsB was flown in **(Supplementary Figure 2F-H)**. We argued that these transient fluorescence intensity increases could be the result of transient pMcsB binding events to the CtsR-DNA complex, which induced a stronger PIFE effect and was reflected in the up-shifts observed in the intensity histograms **(Figure 2A)**. For clarity, the phenomenon was herein referred to as “turbo” PIFE **(Figure 2C)**. Control experiments showed that the “turbo” PIFE effect was not due to direct interactions between pMcsB and the target DNA **(Supplementary Figure 2I)**.

To further validate the interaction between pMcsB and the DNA-bound CtsR, we made a HaloTagged McsB protein and labeled it with the Abberior STAR RED (AS RED) fluorophore. As determined by the ability to inactivate the CtsR-DNA binding, pMcsB-HaloTag was kinase-active but showed a decreased activity as compared to the untagged wild-type kinase **(Supplementary Figure 3A)**. Specific binding events of the pMcsB-HaloTag to the DNA-bound CtsR were observed, which were absent when only the non-specifically bound CtsR, if any, or the target DNA alone was on the imaging surface **(Figure 3A)**. To measure the binding and dissociation rates, we performed single-molecule real-time binding assays^37^ on pMcsB-HaloTag with the surface-tethered CtsR-DNA complex **(Figure 3B)**. Following the flow-in of pMcsB-HaloTag, it began to bind transiently and repeatedly to the same locations (with a radius < 40 nm), indicating specific and transient binding of pMcsB to the same CtsR-DNA complexes. Based on the fluorescence intensity trajectories we obtained, the *k_on_* and *k_off_* rates of pMcsB-HaloTag were estimated to be 0.75 μM^-1^ s^-1^ and 0.34 s^-1^, respectively **(Supplementary Figure 3B-C and Methods)**. The frequency of pMcsB-HaloTag binding events observed here appeared to be lower than that observed with the “turbo” PIFE effect **(Figure 2B and Supplementary Figure 2G)**, possibly due to 1) that the HaloTag variant had a diminished affinity and 2) that the pMcsB-HaloTag concentration was limited (*i.e.*, 15 nM *vs.* 200 nM; >50 nM fluorescently labeled species would generate a strong fluorescence background that interfered with single-molecule detection). The *k_off_* rate measured here is similar to that measured with the “turbo” PIFE effect (0.29 s^-1^, **Figure 2B)**. The *K_D_* calculated from these values is between 387 nM (“turbo” PIFE) and 453 nM (single-molecule binding assays), close to that measured between *B. subtilis* CtsR and McsB.^18^

**Figure 3.**
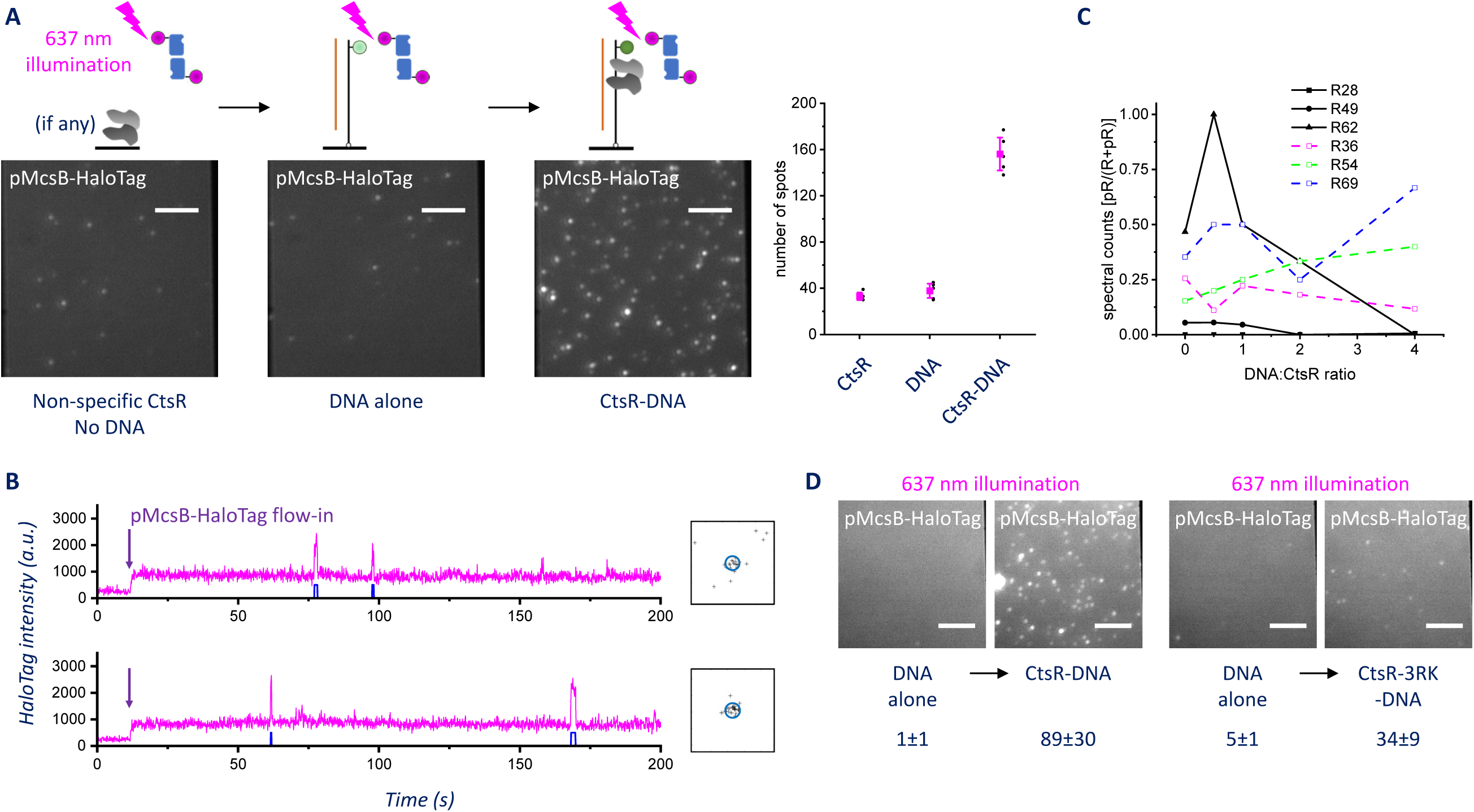
Single-molecule analysis of the pMcsB-HaloTag-CtsR-DNA interaction. (A) Fluorescence images of the AS RED-labeled pMcsB-HaloTag interacting with non-specifically bound CtsR (left), the DNA probe alone (middle) and the CtsR-DNA complex (right). The images were displayed at the same contrast with ImageJ. The scale bars correspond to 5 μm. The average spot counts of pMcsB-HaloTag over an area of 1250 μm^2^ were calculated. The black arrows indicate experiments that were sequentially performed in the same flow channel. 50 nM CtsR was incubated in the flow channel for 5 min and washed away before 30 nM pMcsB-HaloTag was added and imaged. Following this, 10 nM DNA probe was incubated for 2 min and washed away before 30 nM pMcsB-HaloTag was added and imaged. After this, 50 nM CtsR was incubated for 5 min (now in the presence of the immobilized DNA probe from the previous step) and washed away before 30 nM pMcsB-HaloTag was added and imaged. 5 random areas were imaged to calculate the pMcsB-HaloTag spot counts. (B) Example fluorescence intensity trajectories of real-time pMcsB-HaloTag binding to the CtsR-DNA complex upon the addition of 15 nM pMcsB-HaloTag and 0.5 mM ATP in the flow cell. The McsB-bound and unbound dwell times were extracted from the trajectories. The black “plus” marks shown on the right indicate the localization events in the vicinity of the corresponding single molecule. The blue circle has a radius of 40 nm. Only those belong to the same cluster (inside the circle) were included in the dwell time analysis. (C) The relative spectral counts (phosphorylated peptides/total peptides) for positions 28, 49, 62, 36, 54 and 69 at different DNA:CtsR ratios at 42°C. (D) Fluorescence images of the AS RED-labeled pMcsB-HaloTag interacting with the wild-type CtsR-DNA complex (left) and the mutant CtsR-3RK-DNA complex (right). The images were displayed at the same contrast with ImageJ. The scale bars correspond to 5 μm. The average spot counts (±standard deviation) of pMcsB-HaloTag over an area of 1250 μm^2^ were indicated under the respective images. The black arrows indicate experiments that were sequentially performed in the same flow channel. 10 nM DNA probe was incubated for 2 min and washed away before 15 nM pMcsB-HaloTag was added and imaged. After this, 25 nM CtsR or CtsR-3RK was incubated for 5 min (now in the presence of the immobilized DNA probe from the previous step) and washed away before 15 nM pMcsB-HaloTag was added and imaged. 5-10 random areas were imaged to calculate the pMcsB-HaloTag spot counts.

In previous mass spectrometry (MS) and protein mutant studies, 8 arginine residues on CtsR can be phosphorylated by McsB.^9^ Among these 8 residues, 3 arginine residues establish substantive interactions with the DNA major and minor grooves (R28, R49 and R62, *i.e.*, the core arginine residues).^9^ Phosphorylation or phospho-mimic mutations of these key arginine residues, especially R62, completely abolishes the DNA binding activity of CtsR.^9^ In our results, CtsR stayed bound to the DNA during its interaction with pMcsB **(Figure 2)**, indicating the interactions between the core arginine residues and the DNA were largely unaffected. To determine which arginine residues are phosphorylated by pMcsB upon its interaction with the CtsR-DNA complex, we conducted liquid chromatography (LC)-MS/MS experiments. Briefly, CtsR without and with different amounts of DNA were reacted with pMcsB and ATP at 42°C. The reaction solutions were processed and digested to generated the tryptic peptides for LC-MS/MS analysis. For the CtsR sample without DNA, we were able to detect peptides containing phosphorylated arginine residues at positions 49, 62, 36, 54 and 69. However, with increasing amounts of DNA, the relative spectral counts for phosphorylated peptides containing R49 and R62 decreased and became undetectable at the highest DNA:CtsR ratios, whereas the relative spectral counts for phospho-R36, R54 and R69-containing peptides were largely unchanged **(Figure 3C)**. This observation indicates that the periphery arginine residues can be phosphorylated by McsB in the presence of DNA, and is in agreement with the previous results that the DNA-bound CtsR exhibits a reduced rate of phosphorylation at the core arginine residues by McsB, possibly because these residues are deeply buried and protected from the kinase by the DNA backbones.

Based on the MS results, we therefore hypothesize that pMcsB could interact with the DNA-bound CtsR through the periphery arginine residues that are exposed to solution. To test this hypothesis, a CtsR-3RK mutant was made and tested in which three periphery arginine residues (R36, R54 and R69) were mutated to lysine. CtsR-3RK bound to the DNA target with a moderately reduced affinity, possibly due to its slightly perturbed structure upon mutation **(Supplementary Table 1A)**, but the majority of CtsR-3RK remained bound to the DNA 5-10 min after the excess protein in solution was washed away **(Supplementary Figure 3D)**. In comparison with the wild-type CtsR, the 3RK mutations decreased the interaction between the DNA-bound CtsR and pMcsB-HaloTag **(Figure 3D)**. Although we cannot rule out that the structural perturbations caused by the mutations played a role here, this result suggests that the CtsR-McsB interaction observed above is partly mediated through the periphery arginine residues.

Previous studies demonstrate that the glycine-rich loop in CtsR acts as a temperature sensor and regulates the CtsR-DNA binding affinity in response to temperature changes.^27^ Without the help of McsB, sufficiently elevated temperatures alone can dissociate *L. lactis* and *B. subtilis* CtsR from its target DNA.^27^ However, for *G. stearothermophilus* CtsR (the same species examined in this study), the CtsR-DNA complex remains stable even at 55°C, highlighting its extremely tight binding affinity.^27^ We then explored the potential effects on the thermostability of the CtsR-DNA complex caused by its interaction with pMcsB. To test this in our experiments, we flowed in a heated buffer in the flow cell, which subjected the CtsR-DNA complex to elevated temperatures and quickly washed away the dissociated CtsR to prevent rebinding **(Supplementary Figure 4A)**. The CtsR-DNA complex was stable and tolerated pulses of heated buffer washes up to 42°C **(Figure 4A)**. We wonder if treating the CtsR-DNA complex with pMcsB could alter its tolerance to elevated temperatures. After pre-treating the DNA-bound CtsR with pMcsB and ATP for 10 min, CtsR became susceptible to heated buffer washes at 42°C **(Figure 4B)**. We tried to exclude pMcsB and ATP in the heated washing buffer, with the reason that including pMcsB in the heated washing buffer could introduce additional effects that only occurred between CtsR and pMcsB at the higher temperatures. The results showed that even when pMcsB was removed prior to the temperature jump and excluded from the heated washing buffer, the pre-treated CtsR was still heat-susceptible at 42°C **(Figure 4C)**. To test if this effect was related to phosphorylation, the CtsR-DNA complex was treated with McsB in the absence of ATP. Pre-treating the CtsR-DNA complex in the absence of ATP led to a significantly subdued effect of heat sensitization, as heated buffer washes at 42°C induced much less CtsR dissociation **(Figure 4D)**. Therefore, it is likely that the pre-incubation with pMcsB and ATP rendered the DNA-bound CtsR more heat-sensitive through arginine phosphorylation, which may explain why, in our MS experiments conducted at 42°C, the R49 and R62 phosphorylation was detectable at DNA:CtsR ratios close to the stoichiometry (*i.e.*, 0.5:1); only at a large excess, DNA kinetically competed and blocked McsB from phosphorylating R49 and R62 on CtsR.

**Figure 4.**
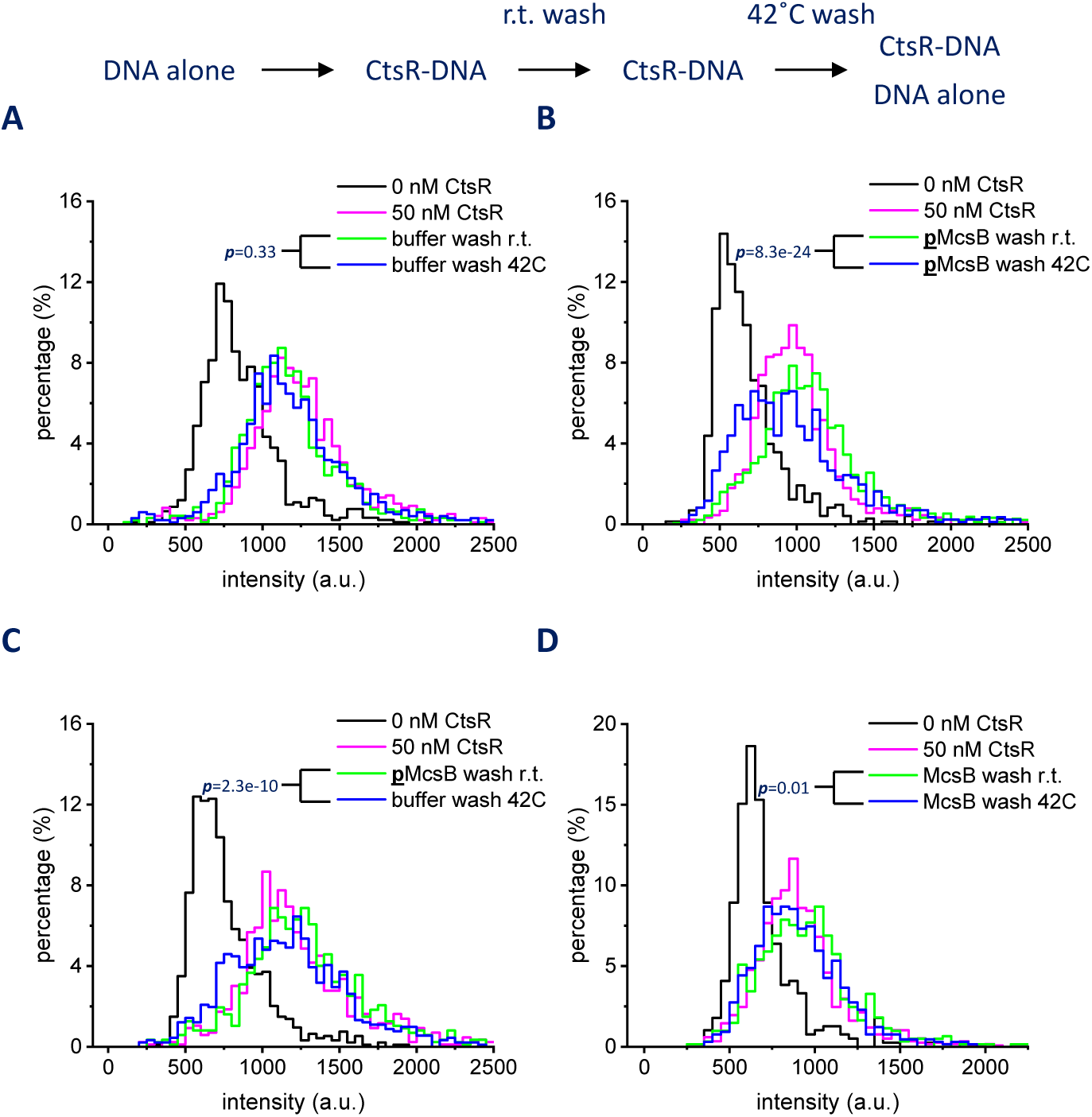
Temperature and phosphorylation-induced CtsR dissociation. (A) Fluorescence intensity histograms of the CtsR-DNA complex before and after a buffer wash at 42°C. The complex was incubated in the buffer for 10 min at room temperature before the heated wash. (B) Fluorescence intensity histograms of the CtsR-DNA complex before and after a pMcsB and ATP wash at 42°C. The complex was first treated with pMcsB and ATP for 10 min at room temperature before the heated wash. (C) Fluorescence intensity histograms of the CtsR-DNA complex before and after a buffer wash (without pMcsB and ATP) at 42°C. The complex was first treated with pMcsB and ATP for 10 min at room temperature before the heated wash. (D) Fluorescence intensity histograms of the CtsR-DNA complex before and after a McsB wash (without ATP) at 42°C. The complex was first treated with McsB in the absence of ATP for 10 min at room temperature before the heated wash. In (A-D), the asymptotic *p*-values are from the two-sample Kolmogorov-Smirnov test. r.t. stands for room temperature. The mean intensity of each selected single molecule was pooled for histogram construction **(Methods)**. Within each group of the intensity histograms, the CtsR addition and washing steps were sequentially performed in the same flow channel. 5-10 random areas containing 919±308 (mean±standard deviation) single molecules were imaged to construct each intensity histogram.

To understand how periphery arginine phosphorylation could make CtsR more sensitive to heat, we used AlphaFold3^38^ to predict the structure of the complex formed between CtsR and the target DNA. Because arginine phosphorylation is not available in the model of AlphaFold3, we chose the arginine-to-aspartic acid mutation as a mimic of arginine phosphorylation. All the mutants we examined, including the CtsR-3RK mutant mentioned above, displayed a RMSD of less than 0.5 Å as compared to the wild-type CtsR structure, suggesting that the point mutations had a minimal effect on protein folding **(Supplementary Figure 4B-C and Supplementary Table 1A)**. To estimate and compare the binding strength of the various CtsR-DNA complexes, we characterized the hydrogen bonds formed between the DNA and CtsR or its phospho-mimic mutants **(Supplementary Table 1B)**. For instance, the wild-type CtsR formed 21 pairs of hydrogen bonds with the DNA. The triple phospho-mimic mutant of the three core arginine residues (R28, R49 and R62) led to a drastic weakening in hydrogen bonding, forming only 15 pairs, whereas mutations of the three periphery arginine residues (R36, R54 and R69) only modestly diminished the strength of the complex, forming 21 pairs of hydrogen bonds but with a slightly longer average bond length **(Supplementary Figure 4D and Supplementary Table 1B)**. The other 6 possible single and double phospho-mimic mutants of R36, R54 and R69 were also examined, all of which were observed to form 21 pairs of hydrogen bonds with almost identical average bond length to the wild-type complex.

Since predicting protein folding and interactions with the phospho-mimic mutations may not fully recapitulate what happens upon adding phosphate groups on a folded structure (*e.g.*, due to kinetic reasons), we performed all-atom molecular dynamics simulations using GROMACS to further elucidate at the amino acid level how periphery arginine phosphorylation modulates the strength of the CtsR-DNA interaction. To assess the compactness of the CtsR-DNA complex, we measured the radius of gyration (RG). The RG of the unphosphorylated CtsR-DNA complex exhibited only minor fluctuations, indicating a stable structure. When the temperature increased from 298K to 323K, the RG did not show a consistent change **(Figure 5A)**, suggesting the complex is not thermosensitive, consistent with our experimental observation above. However, in simulations with the phosphorylated complex at three peripheral arginine residues (*i.e.*, R36, R54, and R69), the RG increased more noticeably with increasing temperatures **(Figure 5B)**, indicating that the complex becomes less compact and exhibits a thermosensitive behavior. We then directly calculated the binding free energy of the CtsR-DNA complex using Molecular Mechanics Poisson-Boltzmann Surface Area (MMPBSA) analysis **(Figure 5C)**. The results showed that the phosphorylated CtsR binds DNA with a significantly higher free energy than the unphosphorylated one, indicating a weaker interaction between phosphorylated CtsR and DNA. Further energy component analysis **(Supplementary Figure 5)** revealed that phosphorylation greatly reduces the electrostatic interactions of the CtsR-DNA complex, consistent with the introduction of additional negative charges by arginine phosphorylation. To further explore this effect, we analyzed the Root Mean Square Fluctuation (RMSF) of the CtsR residues, comparing the phosphorylated and unphosphorylated states (ΔRMSF = RMSF_phosphorylated_−RMSF_unphosphorylated_, **Figure 5D**). Phosphorylation altered the flexibility of specific residues, especially two clusters between I23–I31 and G66–K76. At 298K, both clusters exhibited negative ΔRMSF values, suggesting a decreased flexibility in the phosphorylated CtsR. However, with increasing temperatures, the I23– I31 cluster’s negative ΔRMSF values diminished, while the G66–K76 cluster showed positive ΔRMSF values, indicating a temperature-dependent phosphorylation effect on the CtsR residue flexibility. Notably, both residue clusters contain phosphorylated arginine residues (*i.e.*, R28 and R69, respectively) and are in proximity to DNA **(Figure 5E-F)**, suggesting that phosphorylation at peripheral arginine residues affects neighboring residue dynamics and potentially the CtsR-DNA interaction.

**Figure 5.**
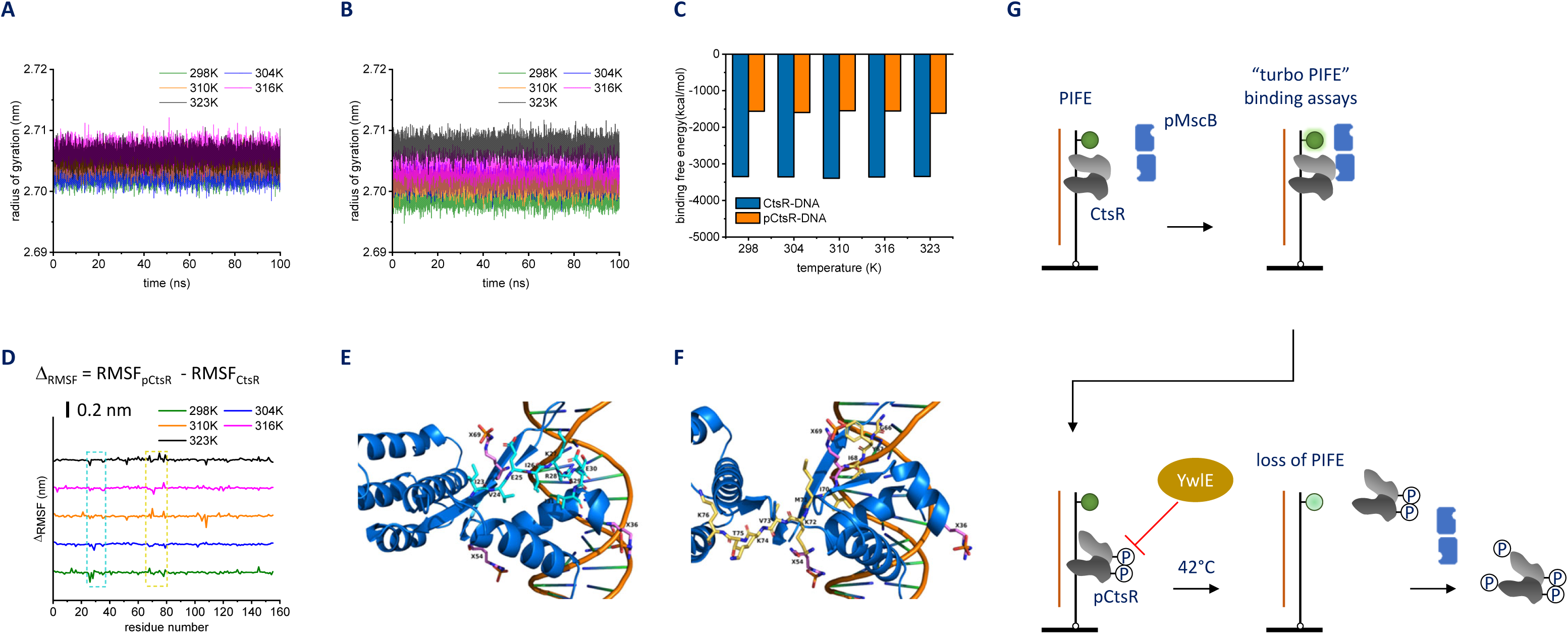
Molecular dynamics simulations on the CtsR-DNA complexes. (A) Radius of gyration (RG) of the unphosphorylated CtsR-DNA complex. (B) RG of the phosphorylated CtsR-DNA complex at three peripheral arginine residues (*i.e.*, R36, R54, and R69). (C) Binding free energy of the CtsR-DNA complexes. “pCtsR” denotes the peripherally phosphorylated CtsR. (D) Root Mean Square Fluctuation (RMSF) analysis of the CtsR residues. ΔRMSF was calculated as the RMSF of the phosphorylated CtsR minus that of the unphosphorylated CtsR. Negative ΔRMSF values (sinks) indicate a decreased flexibility upon phosphorylation, while positive values (peaks) indicate an increased flexibility. The cyan dashed box highlights the I23– I31 residue cluster; the yellow dashed box highlights the G66–K76 residue cluster. (E-F) Structure diagrams of the CtsR-DNA complex showing the I23–I31 residue cluster (E) and the G66–K76 residue cluster (F). Phosphorylated arginine residues are labeled as X36, X54 and X69. (G) Schematic showing that the balance of McsB and YwlE influences the phosphorylation states and the heat sensitivity of the CtsR-DNA complex.

## Discussion

Our results show that the CtsR-DNA complex is stable and forms transient interactions with McsB at room temperature. Since McsB is not able to dissociate CtsR from the complex, it is likely that the core arginine residues are protected in the complex and thus not accessible to McsB without a temperature change. In contrast to the core arginine residues, the MS data show that McsB can leave phosphorylation marks on several periphery arginine residues on CtsR, even in the presence of a large excess of the target DNA. Furthermore, through thermostability experiments and mutagenesis analysis of CtsR, we observe the weakening of the CtsR-DNA complex which can be attributed to periphery arginine phosphorylation. In addition, AlphaFold analysis points out a plausible structural basis for this weakening effect by detecting a mild reduction of hydrogen bonding strength with the phospho-mimic mutant, which is further elaborated with our molecular dynamics simulation results. Overall, our findings suggest a regulation mechanism in which McsB actively modifies the phosphorylation states of the DNA-bound CtsR *in situ* in response to outside stimuli **(Figure 5G)**, different from the previous model where McsB captures and inactivates CtsR only after it dissociates from the DNA.

The electrophoretic mobility shift assay (EMSA) is a widely used method to study protein-nucleic acid interactions, in which protein binding is read out from the mass change-induced mobility shift. Here in a similar effort, we monitored the protein binding to nucleic acids based on the spatial proximity change-induced fluorescence intensity shift.^36^ Therefore, we name approach the fluorescence intensity shift assay, FISA. Compared to EMSA, FISA operates at the single-molecule level and in real time, requiring only a minimal sample quantity while providing dynamic information on weak and transient interactions. Since FISA is based on the PIFE effect, it requires a minimum of one excitation and emission channel in the microscope, making the instrument easier to build and more accessible compared to the colocalization or FRET-based assays. Besides, another benefit of the PIFE-based assay is that it can work with unlabeled proteins, circumventing several technical issues including protein labeling, non-specific binding of fluorescently labeled proteins as well as the “concentration barrier” of detecting single-molecule signals when a high concentration of labeled species is present in the solution (*e.g.*, 200 nM McsB, **Figure 2**).^39^ A potential limitation of the PIFE effect is that it requires the protein and fluorophore to be within a short distance range (*e.g.*, within 4 nm).^36^ However, our results demonstrate that, with a proper positioning of the fluorophore, PIFE can be sensitive not only to the presence of direct protein partners (*e.g.*, CtsR, **Figure 1**), but also to indirect and secondary interactions (*e.g.*, McsB, **Figure 2**), showcasing the technique as a versatile tool for studying protein-nucleic acid interactions. In practice, since a protein can exert rather complex effects on the fluorophore, the choice of the fluorophore and its labeling position need to be optimized on a protein-specific basis as outlined in this and previous studies.^40^ To minimize the batch effects coming from variations in illumination conditions (critical for PIFE), we recommend that samples which are directly compared being analyzed in the same flow cell and, if possible, sequentially in the same flow channel, as often conducted in this study.

Based on our observations, we reason that the phosphorylation states of the periphery arginine residues could help integrate inputs from environmental stresses and determine the proper temperature threshold at which cells initiate the heat-shock response. In bacteria, the phosphorylation states of CtsR are combinedly regulated by the kinase McsB (and in complex with McsA *in vivo*) and phosphatase YwlE,^10,41^ which dynamically add and remove phosphorylation marks, respectively. The balance between the enzyme pair is vital to the precise control of the related cellular functions and gene expression regulation. It is known that certain stresses, such as the oxidative stress, inactive the phosphatase YwlE by causing the formation of disulfide bonds within the enzyme,^29^ which results in accumulated phosphorylation modifications on CtsR. In our model, when bacteria are already faced with a mild oxidative stress, the cells might tune its heat tolerance threshold and initiate the heat-shock response at a lower temperature, as a way to protect the cells from extensive damages in a harsh environment of multiple stresses.

## Materials and Methods

### Protein over-expression and purification

The plasmid for *G. stearothermophilus* McsB was a gift of Tim Clausen. The plasmids for *G. stearothermophilus* CtsR and CtsR-3RK, recombinant McsB-HaloTag were synthesized by SYNBIO Technologies. In general, protein over-expression was performed in *Escherichia coli* BL21(DE3) grown in LB-medium supplemented with 50 μg/ml carbenicillin. The cultures for McsB and McsB-HaloTag were grown to an OD600 of 1.0 at 37°C, and 0.5 mM IPTG was added and the culture was shifted to 18°C for overnight expression. The cultures for CtsR and CtsR-3RK were grown to an OD600 of 0.8 at 37°C, followed by induction with 1 mM IPTG for 4 h. Cells were harvested and the pellet was resuspended in lysis buffer (50 mM PBS, pH 7.5 and 150 mM NaCl, 10% glycerol, 1 mM 2-mercaptoethanol, 1 mM PMSF). All proteins were purified by Ni-NTA affinity chromatography using a 1 mL His Trap column (GE Healthcare Life Science). The column was equilibrated with balance buffer (50 mM PBS pH 7.5 and 150 mM NaCl, 10% glycerol, 50 mM imidazole). Bound proteins were eluted in a buffer gradient (Buffer A: 50 mM PBS, pH 7.5, 150 mM NaCl, Buffer B: 50 mM PBS, pH 7.5, 500 mM NaCl, 500 mM imidazole). The proteins were further purified by a Superdex 200 column (GE Healthcare Life Sciences) equilibrated with the balance buffer (50 mM PBS pH 7.5, 150 mM NaCl). Finally, the proteins were concentrated by a 10 kDa Amicon Ultra-0.5 mL centrifugal filter (Millipore) and stored at −80°C for storage.

### Protein labeling

The McsB-HaloTag protein was labeled with the Abberior® STAR RED HaloTag® Ligands (Abberior) following the manufacture’s procedures. The labeled protein was loaded to a column (Beyotime Biotechnology) to remove the excess fluorophore. Labeling efficiency was calculated according to the maximum absorption of the protein and the fluorophore.

### Mutagenesis

Site-directed mutagenesis of *G. stearothermophilus* CtsR expression constructs were performed using the PrimeSTAR HS DNA Polymerase (Takara) following the manufacturer’s procedures. Primers are listed in **Supplementary Table 2**.

### McsB (auto)phosphorylation assays

The McsB or McsB-HaloTag (10 μM) autophosphorylation reaction was performed in 30 mM Tris-HCl pH 7.4, 20 mM KCl, 5 mM MgCl_2_, and 1 mM ATP at 42°C for 5 min.

To phosphorylate CtsR in solution, 200 nM pMcsB or pMcsB-HaloTag was incubated with 50 nM CtsR in the reaction buffer (30 mM Tris-HCl pH 7.4, 20 mM KCl, 5 mM MgCl_2_, and 0.5 mM ATP) at 25°C for 15 min. CtsR phosphorylation was monitored by single-molecule FISA.

### AlphaFold3 simulation

The structure of the CtsR-DNA complex was predicted using the AlphaFold server, with the number of protein copies set to 2. The DNA binding sequence (26 bp) is provided in **Supplementary Table 2**. Random seed values were fixed at 1 for the prediction. Aspartic acid was used as a substitute to mimic the phosphorylated arginine (CtsR-RD) on CtsR. The CtsR-RD-DNA complexes were similarly predicted using the AlphaFold server, employing identical settings to those used for the wild-type complex.

### Molecular dynamics simulations

Molecular dynamics simulations were carried out using GROMACS (version 2024.3) with the amber14sb_OL15 force field.^43^ Each complex was solvated in a cubic box of TIP3P water under periodic boundary conditions, with Na^+^/Cl^-^ ions added for charge neutralization. Energy minimization was conducted using the steepest descent method. Long-range electrostatics were maintained by the Particle-Mesh Ewald (PME) method, with a 1.2 nm cut-off for short-range interactions. Temperature and pressure were controlled using the V-rescale thermostat and Parrinello-Rahman coupling algorithm, respectively. The system was equilibrated with 500 ps simulations in NVT and NPT ensembles. Finally, production simulations were performed for 100 ns at each of the five temperatures, 298K, 304K, 310K, 316K, and 323K, at a timestep of 2.0 fs. Simulation results were extracted using GROMACS utilities. The molecular visualization was performed with PyMOL (version 3.1).

The unphosphorylated CtsR-DNA complex structure was obtained from the Protein Data Bank (PDB ID 3H0D). For the phosphorylated arginine residues, the initial structure was extracted and then modified using GaussView. Geometry optimization and frequency calculations were performed using the density functional theory (DFT) method at the B3LYP/6-311+G(d,p) level in Gaussian 16.^44,45^ The RESP atomic charges were obtained from DFT results using Multifwn.^46^ Customized force field parameters for phosphorylated arginine were added to the GROMACS topology by manually modifying the residue database.

The binding free energy between CtsR and DNA was calculated with gmx_MMPBSA (version 1.5) based on anaconda and AmberTools23 tools.^47^

### Mass spectrometry analysis of arginine residues on CtsR

CtsR (2 μg) was pre-incubated with various amounts of the 26-bp DNA substrate for 5 min in Tris-HCl buffer (pH 8.0) containing 100 mM KCl, 5 mM MgCl_2_ and 1 mM ATP. Subsequently, 2.1 μg of McsB was added to the binding solution, and the mixture was incubated at 42°C for 1 h. The reacted CtsR (600 ng) was treated with 8 M urea at room temperature for 30 min, followed by reduction with 10 mM tris(2-carboxyethyl) phosphine (TCEP, Thermo Fisher Scientific) at 37°C for 40 min. Alkylation was performed with 25 mM iodoacetamide (IAA, Thermo Fisher Scientific) at room temperature for 25 min. The solution was then loaded onto a home-made tip equipped with a 10 kDa filter and centrifuged to remove urea, as well as the excess TCEP and IAA.^48^ The tip was washed three times with 50 mM ammonium bicarbonate (ABC) before trypsin solution (Promega) was added. Digestion was carried out in a 37°C water bath overnight.

For LC-MS Analysis, an UltiMate™ NCS-3500RS nano system was coupled to an Orbitrap Fusion Lumos Tribrid mass spectrometer (Thermo Fisher Scientific) for peptide analysis. Peptides were eluted using a binary gradient of buffer A (0.1% (v/v) formic acid) and buffer B (100% (v/v) acetonitrile) over a 60-minute period. The gradient progressed from 98% buffer A and 2% buffer B to 35% buffer B at a flow rate of 0.35 µL/min. The mass spectrometer was operated in data-dependent acquisition mode, beginning with a full scan (m/z range 375–1800). The Orbitrap resolution for MS/MS scans was set to 15,000, with a cycle time of 2 seconds (isolation width: 1.6 m/z, mass tolerance: 10 ppm). Precursor ions with charge states of 2–7 were selected for fragmentation and placed on a dynamic exclusion list for 30 seconds. The minimum intensity threshold for precursor selection was set to 5 × 10⁴. Fragmentation was performed using higher-energy collisional dissociation (HCD) with a normalized collision energy of 30%. Additionally, the AGC (Automatic Gain Control) target was set to a custom model, and the maximum injection time was configured to 25 ms.

Raw data were analyzed using Proteome Discoverer (version 2.3.0.523, Thermo Scientific) against a combined database consisting of the UniProt Reference Proteome of *E. coli* strain BL21(DE3), common contaminants, and the sequences of *G. stearothermophilus* CtsR and McsB. Methylthio-modification of cysteine was specified as a fixed modification, while phosphorylation of Arg, Lys, His, Ser, Thr, and Tyr, as well as oxidation of Met, were set as dynamic modifications. Due to the impaired tryptic cleavage at phosphorylated arginine, up to three missed cleavage sites were permitted. The peptide mass tolerance was set to 5 ppm, and the fragment ion mass tolerance to 0.02 Da. Quantitative information for CtsR peptide spectrum matches (PSMs) was obtained through spectral counting. At each arginine residue, the spectral counts of phosphorylated and total peptides containing that residue were calculated.

### Bulk fluorescence measurements

The measurements were performed in a Fluorescence Spectrophotometer F-7100 (Hitachi High-Technologies). The samples were excited with a 520 nm and 650 nm wavelength for Cy3 and Cy5, respectively. To measure the fluorescence spectra, a 1X1 cm Quartz cuvette was filled with 2 nM fluorophore-labeled DNA, followed by adding CtsR to the specific concentrations.

### Single-molecule FISA sample preparation

To construct the fluorescently labeled DNA probe, we ordered the DNA oligos with amine modifications from Sangon Biotech., as listed in **Supplementary Table 2**. The NHS ester derivatives of non-sulfonated Cy3, sulfonated Cy3 and sulfonated Cy5 were purchased from Lumiprobe. The oligo labeling reaction was performed in 50 mM HEPES pH 7.4, 150 mM KCl. The excess fluorophores were removed with the oligo clean and concentrator kits (Zymo Research). The oligos were annealed in the same buffer as the labeling buffer.

### Single-molecule imaging system

The single-molecule imaging system was based on an objective-type total internal reflection fluorescence (TIRF) microscope (Nikon, Eclipse Ti-E). To excite the Cy3 and AS RED fluorophores, respectively, a 532-nm laser (Changchun New Industries Optoelectronics Technology, MGL-S-532-B) and a 637-nm laser (Changchun New Industries Optoelectronics Technology, TEM-F-637) were combined with a dichroic mirror (Chroma, ZT543rdc-UF1). The fluorescence emission was collected by an oil immersion objective (Nikon, CFI Apochromat TIRF 100XC) and recorded by a qCMOS camera (Hamamatsu, QRCA-Quest C15550-20UP). Two notch filers (Chroma ZET532nf and ZET635nf) were used to reject the 532-nm and 637-nm excitation lasers, respectively. A motorized xy-stage (Sanying MotionControl Instruments, PMC400) and a z-motor (Prior Scientific, ES10ZE) were used to stabilize and automate the imaging acquisition process.

### Single-molecule FISA data acquisition and analysis

The biotinylated Cy3-labeled DNA probe was immobilized on a passivated glass coverslip via the surface-tethered biotin and NeutrAvidin. CtsR, CtsR-3RK, McsB and its variants was dissolved at the specified concentrations in the binding buffer (30 mM Tris-HCl pH 7.4, 20 mM KCl, 5 mM MgCl_2_, 10 mM β-mercaptoethanol, 2% glycerol, 4 mM Trolox, 0.8% (w/v) glucose, glucose oxidase and catalase with or without 0.5 mM ATP) and added manually to the flow channel.^9,49^ A volume of 100 μl buffer was added to the flow channel continuously in one go, lasting about 1-2 seconds. Considering a flow channel volume of 10 μl, this 10X volume added could ensure a complete buffer exchange. For the flow-in experiments, a syringe-and-reservoir system was used to manually draw buffers through the flow channel, at a similar flow rate of about 50 μl per second. The reservoir was directly glued to the flow channel entrance to avoid a “dead volume” in the system. In Figure 4, a syringe pump was also tried in addition to the manual drawing method. The tubing connecting the heated reservoir to the flow channel entrance had a volume of 30 μl (calculated from the inner diameter and length of the tubing), thus introducing a “dead volume” in the system. To minimize the variation in the illumination, DNA density, surface passivation efficacy and *etc.*, a group of experimental conditions, if possible, were sequentially imaged in the same flow channel of a flow cell.

For each single-molecule movie, a minimum of 10 frames were taken at the specified frame rate. For background subtraction, a background image for each single-molecule movie was generated with a 16×16-pixel tiling-window median value, interpolation and smoothing algorithm. For single molecule selection, a 10-frame averaged image was generated for each single-molecule movie. Single-molecule spots were selected with a local maximum method with three exclusion criteria, *i.e.*, if a spot was 1) too large, 2) too close to another spot, or 3) too close to the edge of the image, it was excluded for analysis. For single-molecule intensity extraction, the 7×7-pixel intensities centered around each selected single molecule were background-subtracted and summed with a Gaussian-weighted method (FWHM ∼3 pixels). Note the movies from the single-molecule real-time binding assays were analyzed with some modifications as described in its own method section. For intensity histograms, unless specified otherwise, each selected single molecule contributed its intensity values from the first 2-9 frames for histogram construction. For other intensity histograms, the mean intensity value from the 2^nd^-9^th^ frames of each single molecule was used for histogram construction. Avoiding the 1^st^ and last frames could prevent a partially illuminated frame due to the shutter opening and closing. The experimental details were according to refs. ^50–52^ and the single-molecule experiment laboratory manual in ref. ^53^. The implementation and parameters of the analysis algorithms were adapted from refs. ^50–52^. The custom made MatLab and Python codes are available upon request.

To alleviate photobleaching and blinking of fluorophores, the imaging buffer (buffer containing 4 mM Trolox, 0.8% (w/v) glucose, 165 U/ml glucose oxidase and 2170 U/ml catalase) was prepared according to ref. ^49^.

The passivated imaging surfaces used in this study were prepared according to ref. ^50^.

### Single-molecule real-time binding assays

A syringe-and-reservoir system was constructed for the flow-in experiments. The biotinylated Cy3-labeled DNA probe (100 pM) was immobilized on a passivated glass coverslip via the surface-tethered biotin and NeutrAvidin. 25 nM CtsR was diluted in the binding buffer as described above and added manually to the flow channel. pMcsB-HaloTag was added to the reservoir prior to the start of movie acquisition. Single-molecule movies were taken at 10 frames per second. The buffer in the reservoir containing 15 nM pMcsB-HaloTag was manually drawn into the flow channel shortly after the start of movie acquisition. For xy-drift correction, fluorescent beads were used as fiducial markers.

The single-molecule movies were similarly analyzed as described above, except with an extra super-resolution centroid localization analysis (ThunderSTORM within ImageJ). Note that all specific binding events to the same surface-tethered CtsR-DNA complex should be localized in the same cluster within a radius less than 40 nm. Only single molecules that repeatedly displayed such binding events in the same cluster or that colocalized with a CtsR-DNA spot were selected for dwell time analysis. The McsB-bound and unbound dwell times were extracted to construct histograms, and the kinetic rates were calculated from the average dwell times. The experimental details were according to ref. ^37^. The implementation and parameters of the analysis algorithms were adapted from ref. ^37^. The custom made MatLab and Python codes are available upon request.

To alleviate photobleaching and blinking of fluorophores, the imaging buffer (buffer containing 4 mM Trolox, 0.8% (w/v) glucose, 165 U/ml glucose oxidase and 2170 U/ml catalase) was prepared according to ref. ^49^.

The passivated imaging surfaces used in this study were prepared according to ref. ^50^.

## Author contributions

H.C. and B.H. designed the research; H.C., B.H., and Y.G. performed experiments and analyzed data with the help of X.D. and G.D.; J.H. and Y.F. performed molecular dynamics simulations; H.C., B.H., J.H., Y.F., X.D., G.D., X.X., and Y.Z. wrote the paper; B.H., Y.F., X.X., and Y.Z. funded and supervised the project.

## Acknowledgments

We thank all members of the Boyang Hua group and the Yufen Zhao group, especially Prof. Zhenbin Zhang, Dr. Yang Li and associate Prof. Songsen Fu for their technical support in MS analysis. Besides, we acknowledge the Analysis Center of Institute of Drug Discovery Technology for collecting MS data, and Tim Clausen for gifting the plasmid of *G. stearothermophilus* McsB. This work was supported by the National Natural Science Foundation of China No. 42388101 and No. 92256203 (to Y.Z.), the Fundamental Research Funds for the Central Universities 2024300410 (to B.H.), Ningbo Top Talent Project No. 215-432094250 (to Y.Z.), State Key Laboratory of Analytical Chemistry for Life Science, Nanjing University 5431ZZXM2403 (to B.H.), and the National Natural Science Foundation of China No. 12404241 (to Y.F.).

## Supplementary figure captions

**Supplementary Figure 1.**
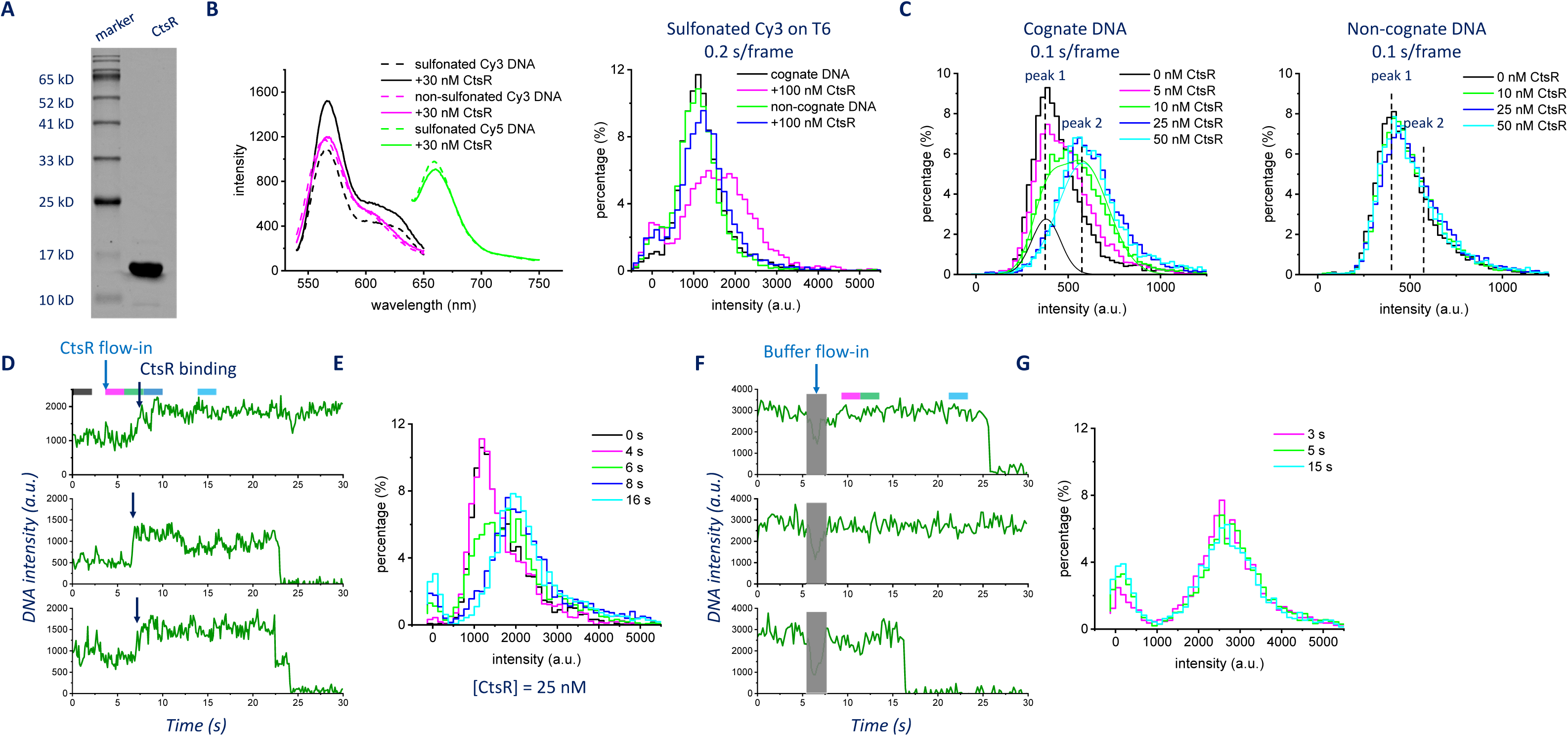
Single-molecule analysis of the CtsR-DNA interaction. (A) SDS-PAGE analysis of the purified recombinant *G. stearothermophilus* CtsR. (B) Bulk fluorescence measurements of the PIFE effects from DNA probes labeled with different fluorophores (left). While sulfonated Cy3 showed a ∼1.5-fold fluorescence enhancement, the non-sulfonated Cy3 and sulfonated Cy5 did not show any PIFE effect. Fluorescence intensity histograms of a DNA probe with sulfonated Cy3 labeled at the T6 position (right). Labeling the fluorophore at T6 (instead of T7) on the DNA weakened CtsR binding, as it was only partially bound by CtsR at 100 nM protein concentration. 5-10 random areas containing 706±171 (mean±standard deviation) single molecules were imaged to construct each intensity histogram. (C) Fluorescence intensity histograms of the cognate (left) and non-cognate (right) DNA probes at different concentrations of CtsR. For the cognate DNA probe at 10 nM CtsR, a multi-Gaussian peak fitting (black and cyan lines) was used to deconvolute the overall peak. We fixed the centers of the two Gaussian peaks at peaks 1 and 2, respectively. However, it was found that the multi-Gaussian peak model (green line) could not fully recapitulate the observed distribution. We cannot rule out a possible middle peak at the intermediate CtsR concentration (*e.g.*, a CtsR monomer instead of dimer binding), especially given the cooperativity we saw with the Hill equation fitting (Figure 1C). For the cognate and non-cognate DNA probes respectively, experiments were sequentially performed in the same flow channel. 5-10 random areas containing 786±76 (mean±standard deviation) single molecules were imaged to construct each intensity histogram. (D) Example fluorescence intensity trajectories of real-time CtsR binding to the target DNA upon the addition of 25 nM CtsR in the flow cell. The frame rate was 0.1 s/frame. The color bars indicate the frames used to construct the intensity histograms of the same color in (E). (E) Time-dependent fluorescence intensity histograms constructed from the acquired trajectories at different time points as indicated by the colored segments in (D). 182 trajectories from the flow-in movie were used to build the histograms. (F) Example fluorescence intensity trajectories of the CtsR-DNA complex upon washing away the excess CtsR in solution. The frame rate was 0.2 s/frame. The color bars indicate the frames used to construct the intensity histograms of the same color in (G). (G) Time-dependent fluorescence intensity histograms constructed from the acquired trajectories at different time points as indicated by the colored segments in (F). 386 trajectories from the flow-in movie were used to build the histograms.

**Supplementary Figure 2.**
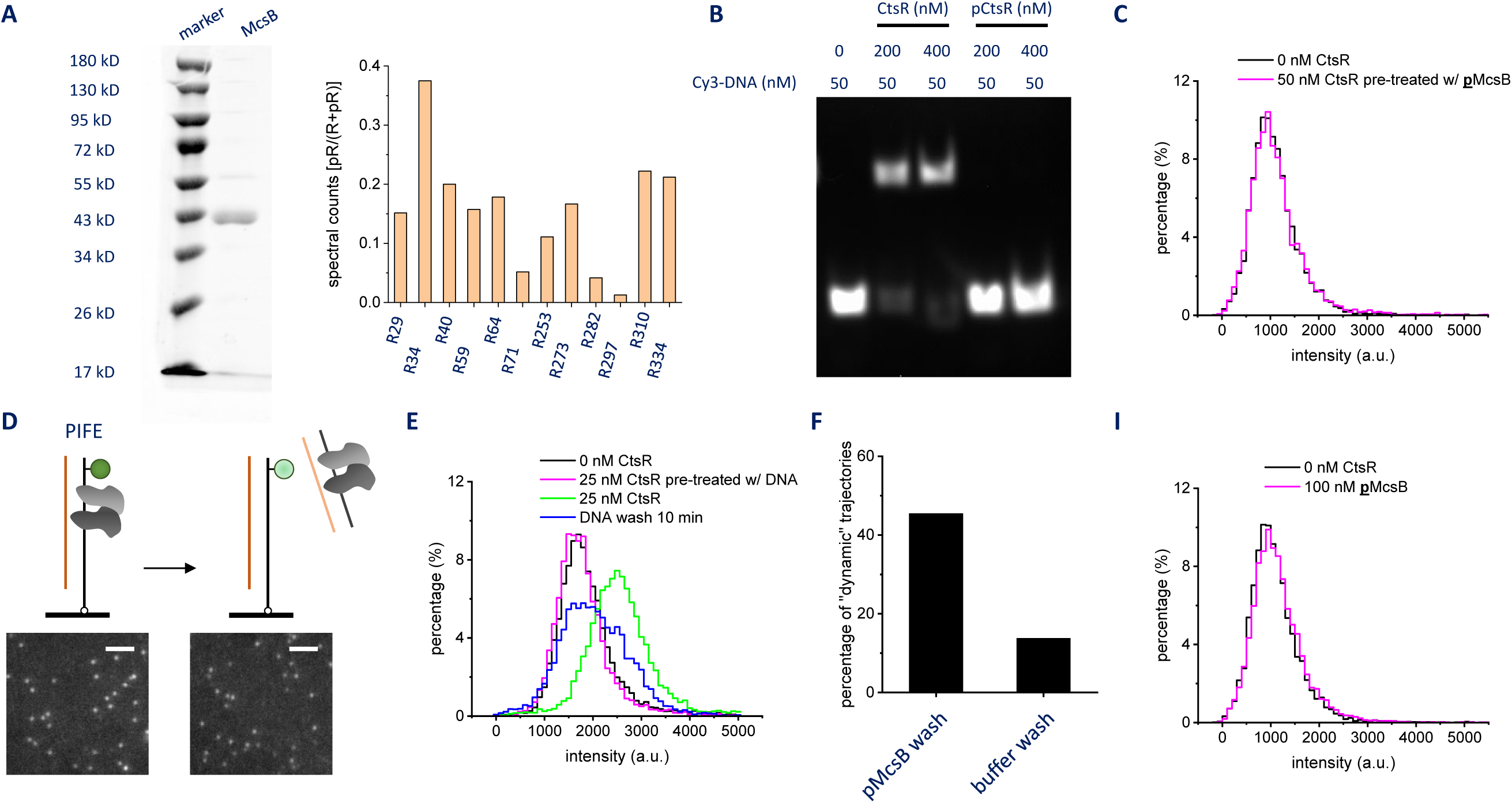
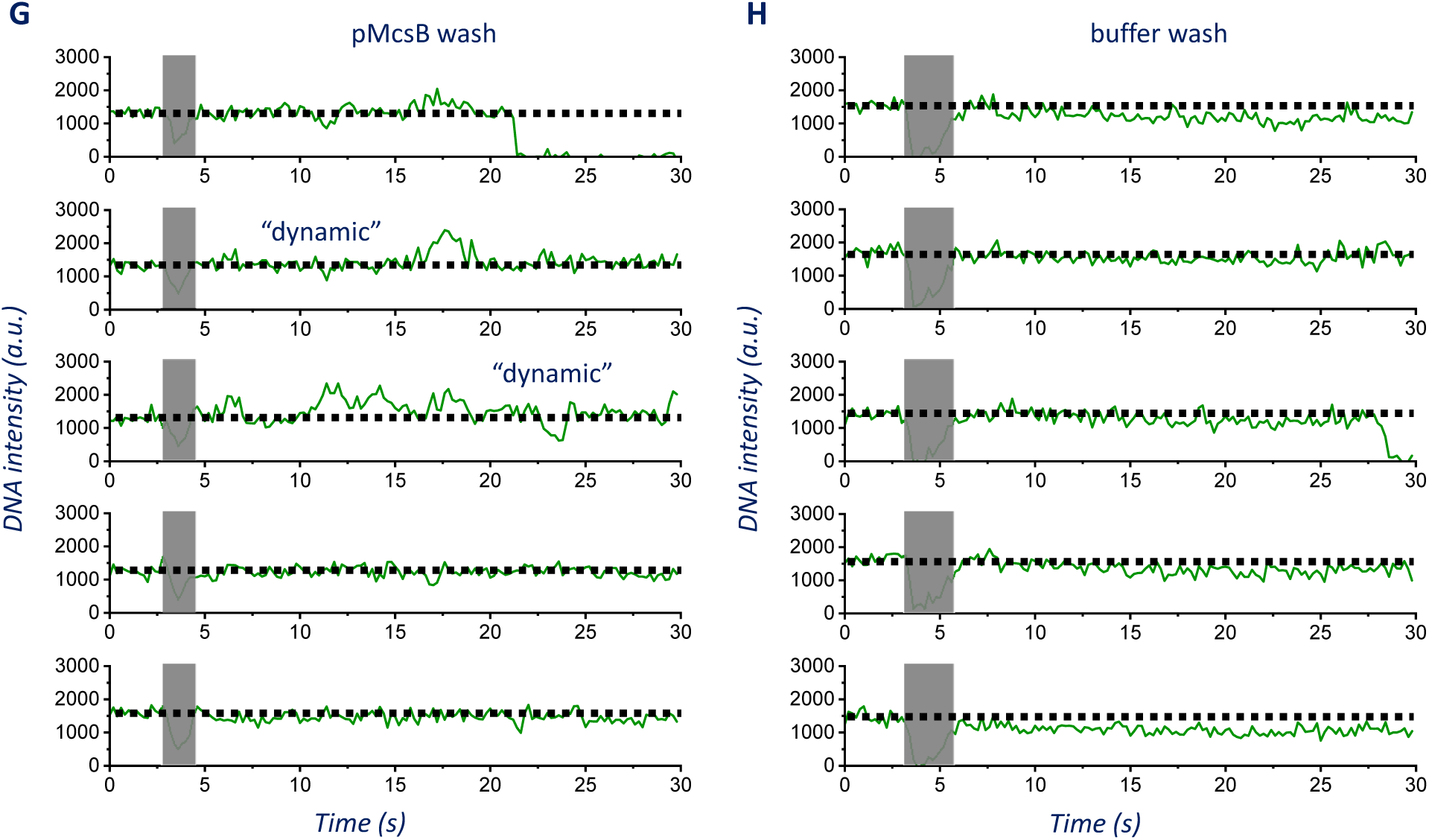
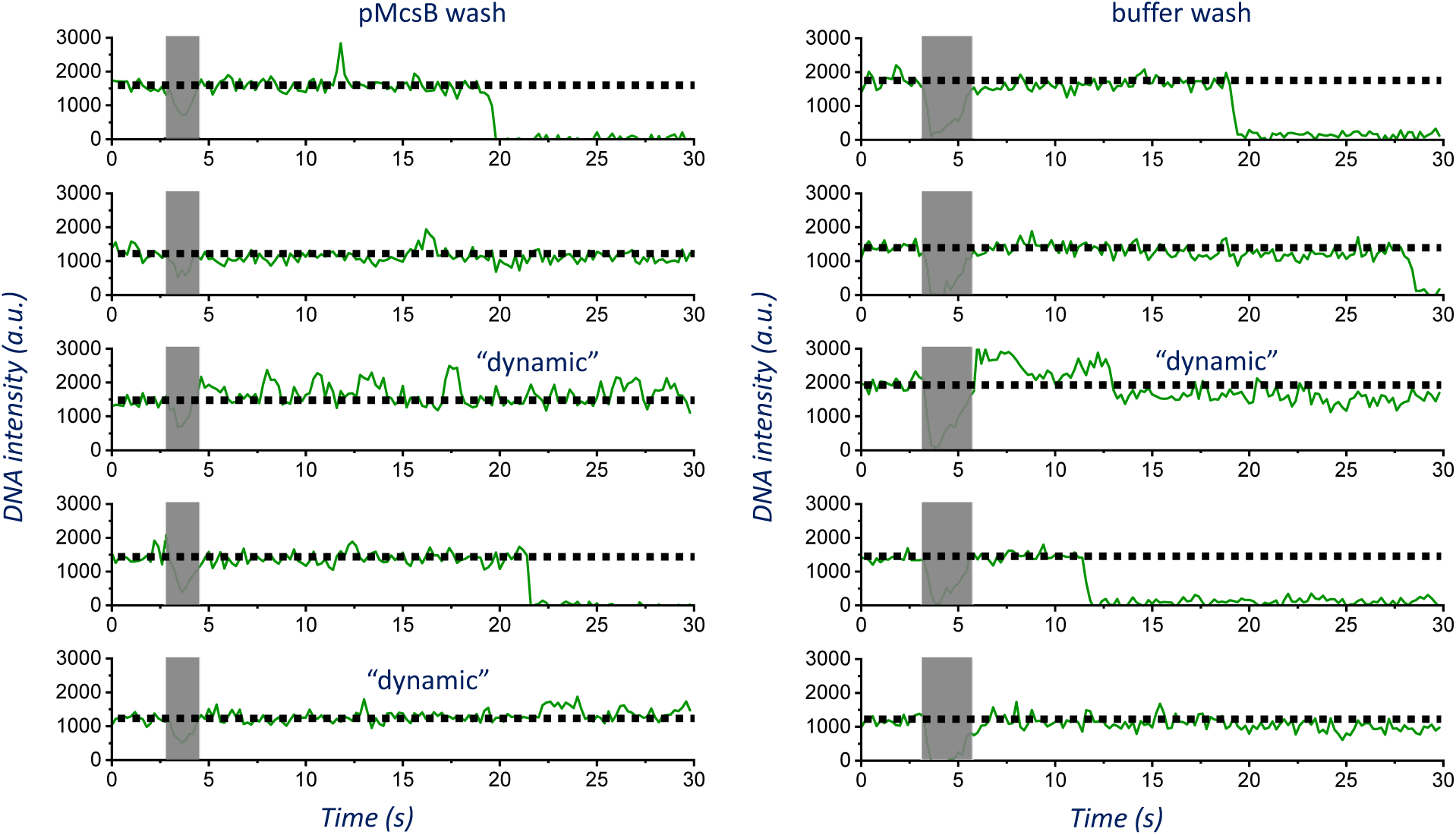
Single-molecule analysis of the pMcsB-CtsR-DNA interaction. (A) SDS-PAGE analysis of the purified recombinant *G. stearothermophilus* McsB (left). The relative spectral counts (phosphorylated peptides/total peptides) at different arginine residue positions on the autophosphorylated pMcsB (right). (B) EMSA analysis of CtsR and McsB activities. pCtsR was the phosphorylated CtsR by pMcsB and ATP before the target DNA was added. (C) Fluorescence intensity histograms of the DNA probe alone (black) and incubated with 50 nM pre-treated CtsR (magenta). CtsR was pre-treated with 200 nM pMcsB and 0.5 mM ATP for 15 min before the protein was added to the flow cell. Experiments were sequentially performed in the same flow channel. 5-10 random areas containing 923 and 753 single molecules were imaged to construct the intensity histograms, respectively. (D) Schematic showing the cognate DNA in solution competing off the bound CtsR as monitored by the loss of PIFE effect. Inserts are representative images of single DNA probes before and after CtsR competing-off. The images were displayed at the same contrast with ImageJ. The scale bars correspond to 2.5 μm. (E) Fluorescence intensity histograms of the DNA probe alone (black), bound by 25 nM CtsR (magenta), after CtsR competing-off (green) and 25 nM CtsR pre-bound with the cognate DNA (blue). When CtsR was pre-bound with the cognate DNA in solution, it cannot bind to the surface-tethered cognate DNA. Experiments were sequentially performed in the same flow channel. 5-10 random areas containing 908±108 (mean±standard deviation) single molecules were imaged to construct each intensity histogram. (F) The percentage of trajectories showing transient spikes after the pMcsB and buffer washes, respectively. 132 and 130 trajectories in the conditions were included in the analysis, respectively. (G-H) Representative fluorescence intensity trajectories of real-time pMcsB binding to the CtsR-DNA complex upon the addition of 200 nM pMcsB and 0.5 mM ATP in the flow cell. The “dynamic” trajectories were manually identified as those displaying a higher intensity level than that of the CtsR-DNA complex (black dotted line). To exclude the intensity noise that occasionally passed the black dotted line, some trajectories were not selected (*e.g.*, trajectories 1 and 7 in G), which might underestimate the percentage of dynamic trajectories. (I) Fluorescence intensity histograms of the DNA probe alone (black) and incubated with 100 nM pMcsB and 0.5 mM ATP (magenta). Experiments were sequentially performed in the same flow channel. 5-10 random areas containing 923 and 847 single molecules were imaged to construct the intensity histograms, respectively.

**Supplementary Figure 3.**
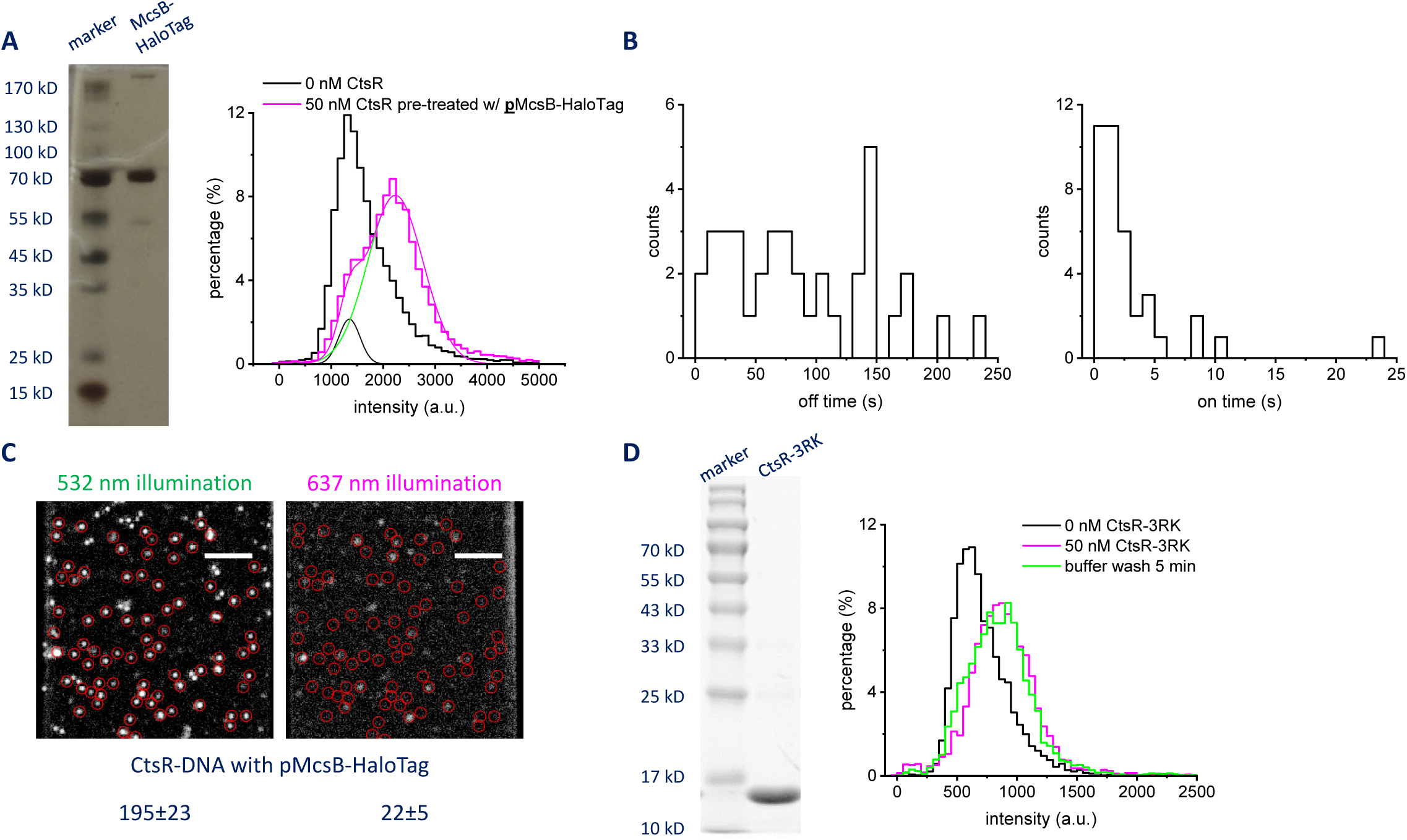
Single-molecule analysis of the pMcsB-HaloTag-CtsR-DNA interaction. (A) SDS-PAGE analysis of the purified recombinant *G. stearothermophilus* McsB-HaloTag (left). Fluorescence intensity histograms (right) of the DNA probe alone (black, step) and incubated with 50 nM pre-treated CtsR (magenta, step). CtsR was pre-treated with 200 nM pMcsB-HaloTag and 0.5 mM ATP for 15 min before the protein was added to the flow cell. Based on a multi-Gaussian peak fitting (magenta line), 91% of the pre-treated CtsR could still bind to the DNA (green line), whereas 9% was phosphorylated and inactivated (black line), indicating the reduced kinase activity of pMcsB-HaloTag. Experiments were sequentially performed in the same flow channel. 5-10 random areas containing 1561 and 2809 single molecules were imaged to construct the intensity histograms, respectively. (B) Dwell time histograms of the pMcsB-HaloTag-unbound durations (T_off_, left) and bound durations (T_on_, right). The dwell times were collected in the first 3 min after the addition of pMcsB-HaloTag in the flow cell. To estimate the *k_on_* and *k_off_* rates, the mean McsB-unbound and bound dwell times were calculated from 38 events from 22 single molecules. (C) Fluorescence images of the AS RED-labeled pMcsB-HaloTag (right) interacting with the wild-type CtsR-DNA complex (left). The images were background-subtracted and displayed at the same contrast with MatLab. The scale bars correspond to 5 μm. The average spot counts (±standard deviation) of the CtsR-DNA complex and pMcsB-HaloTag over an area of 1250 μm^2^ were indicated under the respective images. 100 pM DNA probe, 25 nM CtsR and 12.5 nM pMcsB-HaloTag were sequentially added in the flow channel. 5-10 random areas were imaged to calculate the spot counts. The red circles indicate the selected DNA spots. (D) SDS-PAGE analysis of the purified recombinant *G. stearothermophilus* CtsR-3RK (left). Fluorescence intensity histograms of the mutant CtsR-3RK-DNA complex before and 5 min after a buffer wash to remove the excess CtsR-3RK in solution (right). Experiments were sequentially performed in the same flow channel. 5-10 random areas containing 776, 549, and 498 single molecules were imaged to construct the intensity histograms, respectively.

**Supplementary Figure 4.**
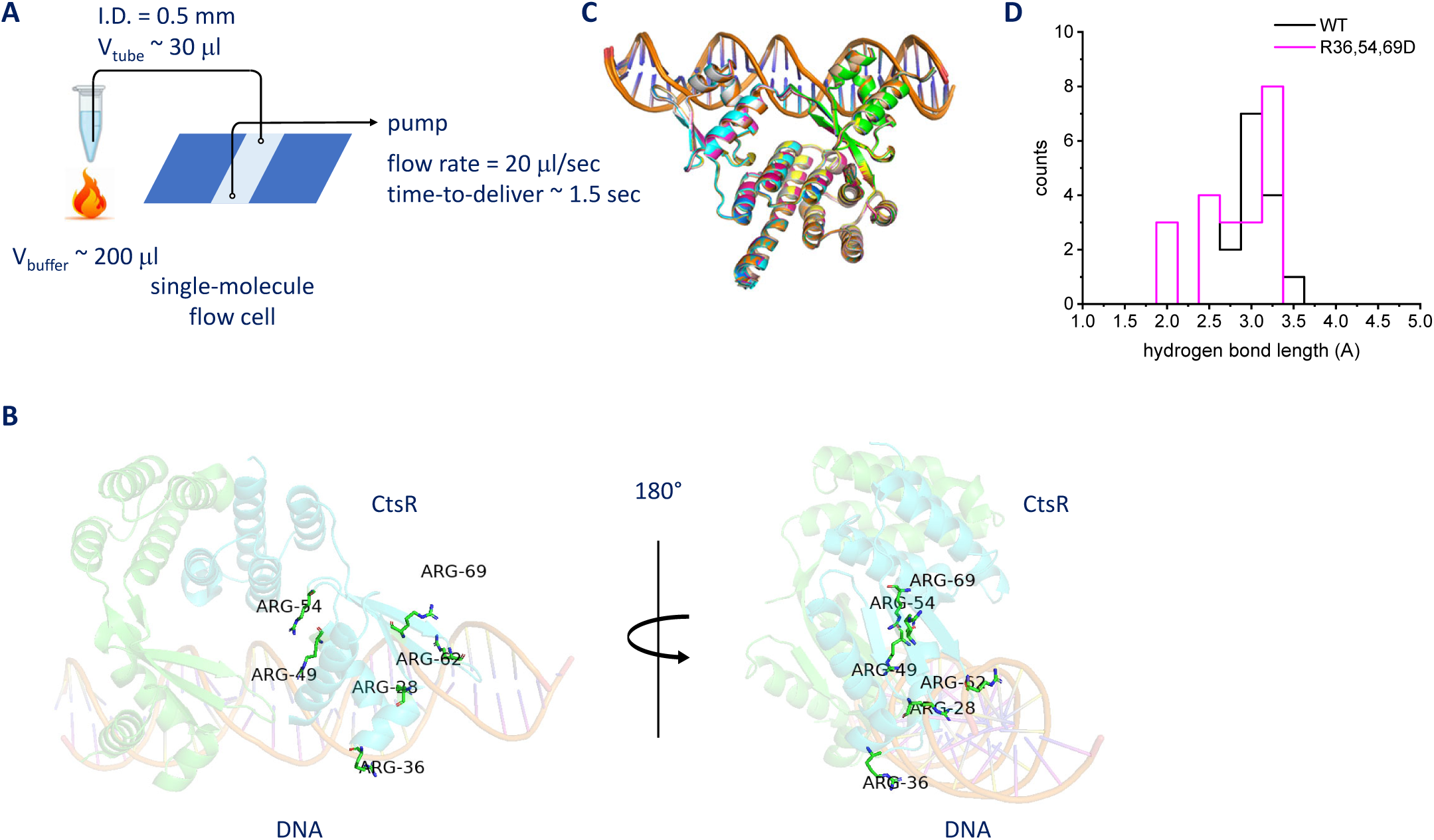
Structural predictions and analysis of the CtsR-DNA complexes with AlphaFold. (A) Schematic showing the device for the heated buffer washes. (B) Structure diagrams of the wild-type CtsR-DNA complex highlighting the core and periphery arginine residues. (C) Structure alignment of the CtsR-DNA complexes from AlphaFold. (D) Hydrogen bond length histograms of the wild-type (black) and R36,54,69D (magenta) CtsR-DNA complexes, respectively.

**Supplementary Figure 5.**
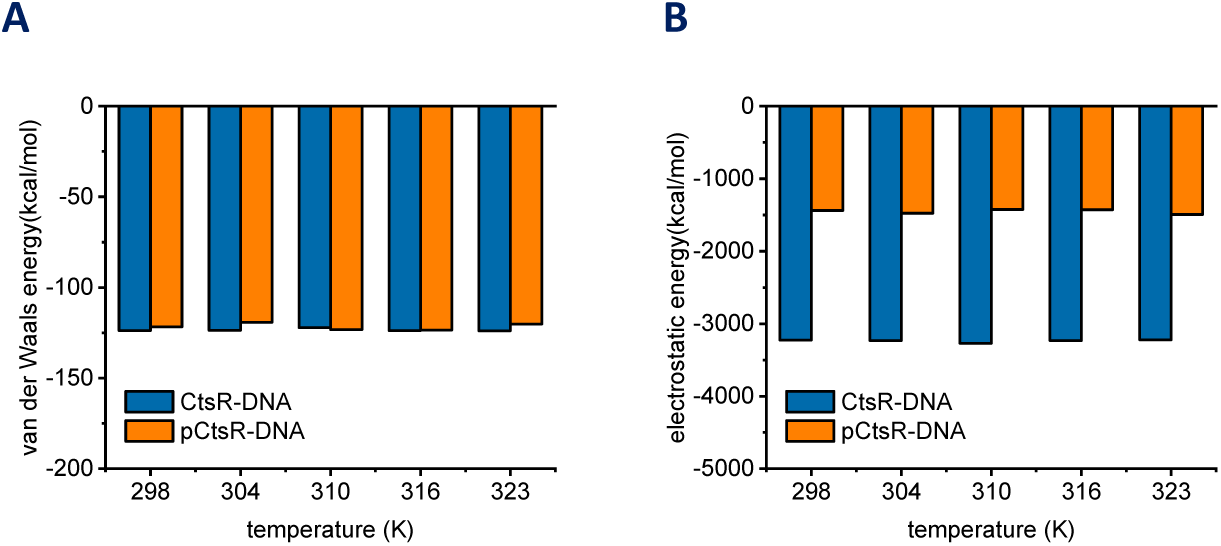
Binding energy analysis of the CtsR-DNA complexes with molecular dynamics simulations. (A-B) The Van der Waals energy (A) and the electrostatic energy (B) of the CtsR-DNA complexes at different temperatures. “pCtsR” denotes the peripherally phosphorylated CtsR.

